# An Active Inference Approach to Dissecting Reasons for Non-Adherence to Antidepressants

**DOI:** 10.1101/743542

**Authors:** Ryan Smith, Sahib Khalsa, Martin Paulus

**Affiliations:** Laureate Institute for Brain Research, Tulsa, OK; University of Tulsa, Department of Community Medicine, Tulsa, OK

**Keywords:** active inference, adherence, psychiatric treatment, computational psychiatry, antidepressants, decision-making, predictive processing

## Abstract

**Background:** Antidepressant medication adherence is among the most important problems in health care worldwide. Interventions designed to increase adherence have largely failed, pointing towards a critical need to better understand the underlying decision-making processes that contribute to adherence. A computational decision-making model that integrates empirical data with a fundamental action selection principle could be pragmatically useful in 1) making individual level predictions about adherence, and 2) providing an explanatory framework that improves our understanding of non-adherence.

**Methods:** Here we formulate a partially observable Markov decision process model based on the active inference framework that can simulate several processes that plausibly influence adherence decisions.

**Results:** Using model simulations of the day-to-day decisions to take a prescribed selective serotonin reuptake inhibitor (SSRI), we show that several distinct parameters in the model can influence adherence decisions in predictable ways. These parameters include differences in *policy depth* (i.e., how far into the future one considers when deciding), *decision uncertainty, beliefs about the predictability (stochasticity) of symptoms*, *beliefs about the magnitude and time course of symptom reductions and side effects*, and the *strength of medication-taking habits* that one has acquired.

**Conclusions:** Clarifying these influential factors will be an important first step toward empirically determining which are contributing to non-adherence to antidepressants in individual patients. The model can also be seamlessly extended to simulate adherence to other medications (by incorporating the known symptom reduction and side effect trajectories of those medications), with the potential promise of identifying which medications may be best suited for different patients.

## Introduction

Medical treatment depends crucially on a patient’s decision to adhere. Unfortunately, the number of patients who follow treatment recommendations is quite low (1). Nearly half of prescribed medications are not taken and roughly 125,000 deaths each year are due to non-adherence; costs associated with non-adherence are between $100 and $300 billion annually (2). In the context of mental healthcare – the focus of this paper – roughly one in five patients adhere to antidepressant medication treatment for more than four months (3), and the majority discontinue within 30 days (4). Low adherence rates do not appear fully attributable to an objective lack of efficacy, as those who follow treatment recommendations show lower recurrence risk (5), cardiovascular mortality (6), overall mortality (7, 8) and lower suicide rates (9). Individuals who adhere also report greater perceived benefits and fewer concerns than those who do not (1). Thus, it is somewhat perplexing that patients often do not adhere.

Beliefs about medication and personality attributes both influence adherence. Adherence to psychiatric interventions is lower in those with lower treatment expectancy (10) and in those who experience sexual side effects (11). Such findings have led to the ‘necessity-concerns framework’ (12) which proposes that adherence decisions involve weighing expected negative outcomes against beliefs about the necessity/efficacy of treatment (13–18). Personality variables associated with greater adherence include higher *persistence* (19, 20), greater *self-efficacy* (21–23), lower *optimism* (24), greater *self-control* (25), and greater *internal locus of control* (26)(27).

Habit formation may also be central to the development of stable health behaviors (28), an effect that could generalize to adherence behavior (29, 30). However, studies have found that the amount of time required to form strong habits is highly variable (e.g., 18 to 254 days in one study (31)). The processes that moderate habit formation time are therefore also relevant to understanding long-term adherence.

While several interventions to promote adherence have been tested, involving educational (32, 33) counseling (34), and coaching (35) approaches, they have shown low efficacy in randomized controlled trials (36). There is therefore a need to better understand the decision-making processes contributing to adherence. A precise and quantitative delineation of these processes could help inform and develop targeted interventions focused on specific adherence-related processes and provide better measures for predicting adherence.

Computational psychiatry approaches have recently gained prominence, due in part to their potential to quantitatively model behavior and illustrate how several underlying processes can contribute to maladaptive perception and decision-making (37–40). In this manuscript we introduce a computational model of antidepressant medication adherence, based on the active inference framework (41), and examine how it might provide a detailed characterization of different probabilistic beliefs and related inference processes that plausibly contribute to adherence decisions. This approach could help clarify the way individual differences in computational processes can arbitrate between adherence and non-adherence in individual patients. We will focus on the initial decision to adhere over the first 12 weeks of treatment, in which symptoms typically decline and stabilize (42) and in which side effects first appear and subsequently reduce somewhat in severity (43). We also simulate processes that may moderate the speed with which individuals develop strong medication-taking habits after their initial decision to adhere.

## Method

### An active inference model of adherence

The active inference framework (41) is one approach to provide a computational framework for adherence. This framework assumes the brain develops a generative model that represents different possible states of the world and generates predictions about the outcomes it will observe if its beliefs are accurate. It uses observations to update the model and to generate sequences of actions (policies) that minimize a statistical quantity called ‘expected free energy’ (**G**); briefly, policies that minimize **G** are those expected to produce the most preferred (rewarding) observations while also maximizing information gain (see table 2). The specific model we use is a Markov decision process (MDP; figure 1), which is particularly useful for modeling decision processes that require consideration of distal future outcomes under uncertainty (44, 45). The formal basis for these models has been thoroughly detailed elsewhere (41, 46–49) (also see tables 1 and 2 for more mathematical detail).

**Figure 1.**
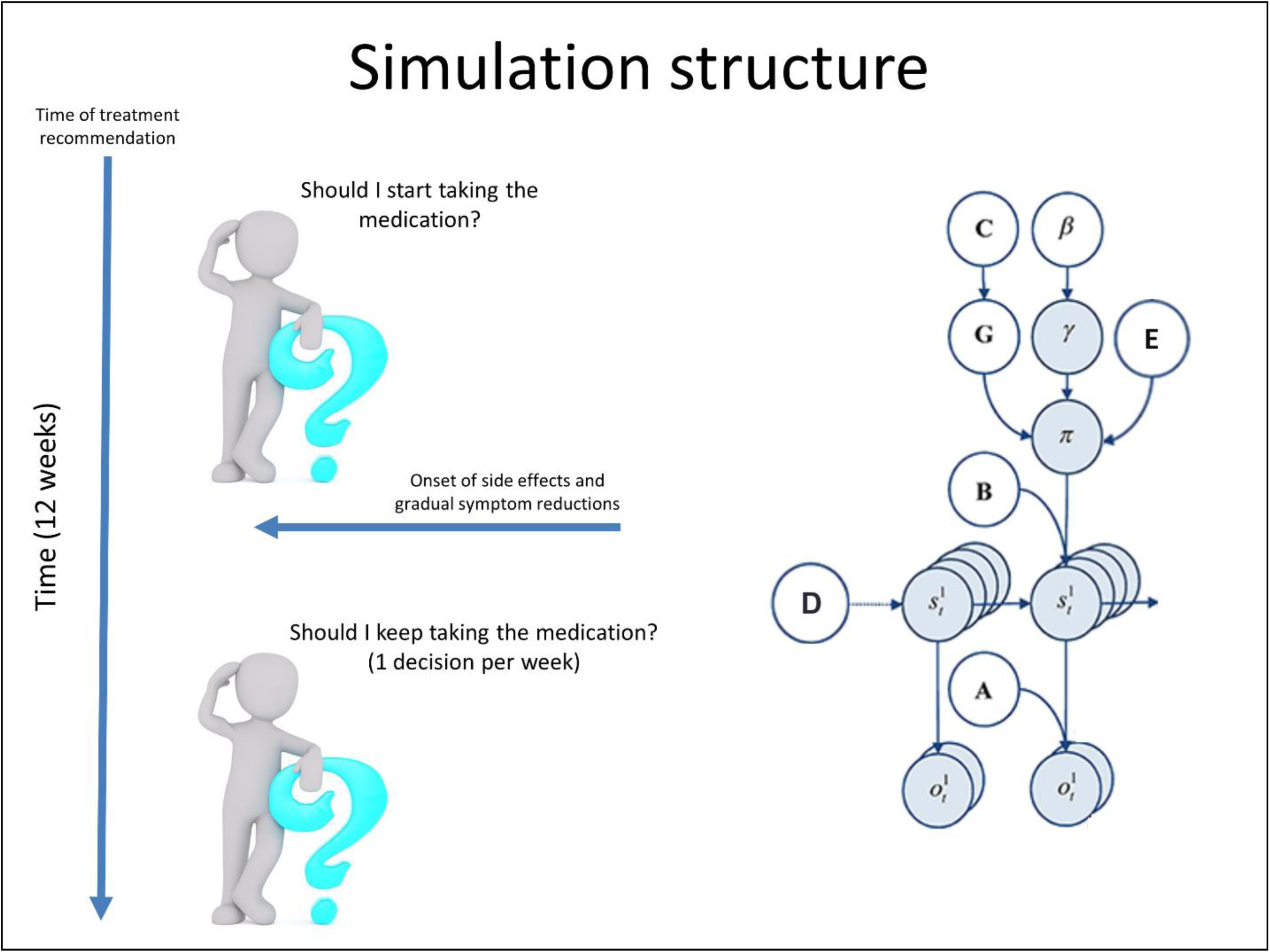
Illustration of the Markov decision process formulation of active inference used in the simulations described in this paper. The generative model is here depicted graphically on the right, such that arrows indicate dependencies between variables. Here observations (*o*) depend on hidden states (*s*), where this relationship is specified by the **A**-matrix, and those states depend on both previous states (as specified by the **B**-matrix, or the initial states specified by the **D**-vector) and the sequences of actions (policies; *π*) selected by the simulated patient. The probability of selecting a particular policy in turn depends on the expected free energy (**G**) of each policy with respect to the prior preferences (**C**) of the simulated patient. The degree to which expected free energy influences policy selection is also modulated by an expected policy precision parameter (**γ**), which is in turn dependent on a prior over expected precision (***β***; the rate parameter of a gamma distribution, *P*(*γ*) = Γ(1, ***β***)) – where higher ***β*** values promote lower confidence in policy selection (i.e., less influence of the differences in expected free energy across policies). The **E**-vector is a prior distribution over policies, p(π), which also influences policy selection and can be thought of as encoding a patient’s habits. For more details regarding the associated mathematics, see (46, 53). In our model, the observations were depression symptom levels and antidepressant side effects, the hidden states included beliefs about progress in treatment over time and beliefs about adherence decisions, and the policies included the choice to adhere or cease adherence at each week over 12 weeks of treatment (e.g., choosing to discontinue on a Wednesday vs. a Saturday of a given week was treated as the same choice in the model; as described in the main text, this modeling choice allowed the integration of week by week empirical data on symptom and side effect trajectories on antidepressants). As depicted on the left, our simulations began at “week 0” when treatment was initially recommended; the simulated patient then chose whether or not to adhere to treatment based on her beliefs about the way that symptoms and side effects would change over time if they did vs. did not adhere (and her subsequent observations over time). Preferences were set such that the patient had stronger and stronger preferences to observe lower and lower symptom levels as well as lower and lower side effect levels.

**Table 1.** Model variables.

| **Model variable** | **General Definition** | **Model-specific specification** |
| --- | --- | --- |
| $O_t$ | Observable outcomes at time $t$ | Outcome modalities:<br>1. Depressive symptom severity (0 - 10) at each week of treatment.<br>2. Side effect severity (0 - 3) at each week of treatment. |
| $S_t$ | Hidden states at time $t$ | Hidden state factors:<br>1. Progress in treatment (0 - 12) at each week of treatment.<br>2. Adherence vs. non-adherence at each week of treatment. |
| $\pi$ | A distribution over action policies encoding the predicted value of each policy. In this model, each policy is a series of allowable actions, where actions correspond to different state transitions (i.e., different '**B**' matrices) that can be chosen by the patient. Policies are chosen by sampling from this distribution. | Allowable policies included the decision to continue taking the medication over 12 weeks or to cease medication at any week prior to week 12. That is, the patient could choose to transition from the state of 'adhering' to the state of 'not adhering' at each week. |
| **A** matrix<br>$p(o_t \vert s_t)$ | A matrix encoding beliefs about the relationship between hidden states and observable outcomes (i.e., the probability that specific outcomes will be observed given specific hidden states). | Encodes beliefs about the relationship between treatment progress, adherence decisions, depressive symptoms, and side effects. |
| **B** matrix<br>$p(s_{t+1} \vert s_t)$ | A matrix encoding beliefs about how hidden states will evolve over time (transition probabilities). For states that are under the control of the patient, there are multiple **B** matrices, where each matrix corresponds to one action (state transition) that the patient may choose at a given time point (if consistent with an allowable policy). | Encodes beliefs about the way treatment progress will change given the choice to adhere or not adhere at each week of treatment. For the hidden state factor corresponding to being in a state of 'adhering' or 'not adhering', distinct **B** matrices encoded the controllable transitions of either remaining in the state of 'adhering' or transitioning to the state of 'not adhering.' |
| **C** matrix<br>$p(o_t)$ | A matrix encoding the degree to which some observed outcomes are preferred over others (technically modeled as prior expectations over outcomes). | Encodes the stronger preference for lower depressive symptoms and lower side effect severities. |
| **D** matrix<br>$p(s_1)$ | A matrix encoding beliefs about (a probability distribution over) initial hidden states. | The simulated patient always begins in an initial state of being “undecided” about choosing to adhere or not, and always begins in an initial state of being at 0 weeks of treatment progress. |
| **E** matrix<br>$p(\pi)$ | A matrix encoding beliefs about what actions will be chosen a priori (a prior probability distribution over policies implemented as a vector assigning one value to each policy), based on the number of times different actions have been chosen in the past. | Higher values in this matrix indicate a greater number of previous choices to adhere vs. not adhere to medications in the past. |

**Table 2.** Model update equations.

| Model update component | Update equation | Model-specific description |
| --- | --- | --- |
| Updating beliefs about initial states expected under each allowable policy | $s_{\pi,1} = \sigma \left( \frac{1}{2} (\ln \mathbf{D} + \ln \mathbf{B} \cdot s_{\pi,2}) + \ln \mathbf{A} \cdot o_t \right)$ | Updating beliefs about initial states (e.g., treatment progress after perceiving initial symptom and side effect levels), based on initial expectations (**D**), initial transition beliefs (**B**), and consistency with new observations ($\mathbf{A} \cdot o$). The $\sigma$ symbol indicates a softmax operation which allows state estimates to make up a proper probability distribution. (Note: the dot (·) notation here indicates transposed matrix multiplication). |
| Updating beliefs about subsequent states expected | $s_{\pi,t} = \sigma \left( \frac{1}{2} (\ln \mathbf{B} s_{\pi,t-1} + \ln \mathbf{B} \cdot s_{\pi,t+1}) + \ln \mathbf{A} \cdot o_t \right)$ | Updating beliefs about subsequent states over time (e.g., treatment progress over time based on observed |
| under each allowable policy over time | | symptom and side effect levels at each subsequent week of treatment), based on prior beliefs about transition probabilities (**B**) associated with each possible policy and consistency with new observations (**A** · **o**). The $\sigma$ symbol indicates a softmax operation which allows state estimates to make up a proper probability distribution. (Note: the dot (·) notation here indicates transposed matrix multiplication). |
| Probability of selecting each allowable policy | $\pi = \sigma(\ln \mathbf{E} - \gamma \mathbf{G} - \mathbf{F})$ | The distribution over policies from which actions are sampled (e.g., encoding the probability that the patient will choose to adhere or not adhere). This is determined by learned priors over policies (**E**), the expected free energy of each allowable policy (**G**), and the observed free energy (**F**) of each policy with new observations over time. The influence of expected free energy is modulated by an expected policy precision term ($\gamma$) encoding the expected uncertainty of desired outcomes. The $\sigma$ symbol indicates a softmax operation which allows policy values to make up a proper probability distribution. |
| Expected free energy of each allowable policy | $\mathbf{G}_{\pi} = \sum_t (\mathbf{o}_{\pi,t} \cdot (\ln \mathbf{o}_{\pi,t} - \ln \mathbf{C}) - \text{diag}(\mathbf{A} \cdot \ln \mathbf{A}) \cdot \mathbf{s}_{\pi,t})$ | This quantity evaluates the value of each policy based on beliefs about the ability of each policy to lead to the most desired outcomes (e.g., the lowest combination of symptoms and side effects over time), while also reducing uncertainty about states. Achieving the most desired outcomes corresponds to minimizing the divergence between the observations expected conditional on each policy ($\mathbf{o}_{\pi,t}$) and preferred observations (**C**). (Note: when applied to matrices, the dot (·) notation here indicates transposed matrix multiplication). |
| Free energy of each allowable policy | $\mathbf{F}_\pi = \sum_t \mathbf{s}_{\pi,t} \cdot \left( \ln \mathbf{s}_{\pi,t} - \frac{1}{2} (\ln \mathbf{B} \mathbf{s}_{\pi,t-1} + \ln \mathbf{B} \cdot \mathbf{s}_{\pi,t+1}) - \ln \mathbf{A} \cdot \mathbf{o}_t \right)$ | This quantity evaluates the evidence that inferred states provide for each policy based on new observations at each time point. In this case, inferred changes in treatment progress over time provide evidence supporting the policy to adhere or not adhere. (Note: when applied to matrices, the dot (·) notation here indicates transposed matrix multiplication). |
| Expected policy precision | $P(\gamma) = \Gamma(1, \beta)$ $\beta = \beta + (\pi - \pi_0) \cdot \mathbf{G}$ $\gamma = 1/\beta$ | The prior on expected policy precision term ($\beta$) is the 'rate' parameter of a gamma distribution, which is a standard distribution to use as a prior for expected precision ($\gamma$). This latter term modulates the influence of expected free energy on policy selection. When $\beta$ is high (reflecting low confidence about whether one should choose to adhere to medication), policy selection becomes less deterministic and is more strongly influenced by prior habits. When subsequent observations alter the precision of the distribution over policies ($\pi - \pi_0$), the value for $\beta$ is updated to reflect this change in decision uncertainty. |

One approach to examine the computational processes that might affect adherence is to model the decision-making situation for an individual that is newly prescribed an antidepressant for a depressive episode (see figure 2). We selected a time frame of the first 12 weeks since most antidepressant trials focus on this period. Moreover, we utilized previously reported decreases in depressive severity (42) and the temporal emergence of unwanted effects (43). In our model, the policy to adhere or not on (any day during) each of 12 weeks of treatment (13 policies total, formally modeled as one choice per week) was based on initial expectations about how depressive symptom severity (from 0-10) and side effect severity (from 0-3) would change over time given each policy, and on how these expectations were subsequently updated when changes in symptoms and side effects were observed by the simulated patient. The 2 state factors in our model correspond to beliefs about “progress in treatment” (which predict different combinations of observed symptom and side effect levels), and beliefs about whether one is currently choosing to adhere (which predict different patterns of change over time in symptom and side effect levels).

**Figure 2.**
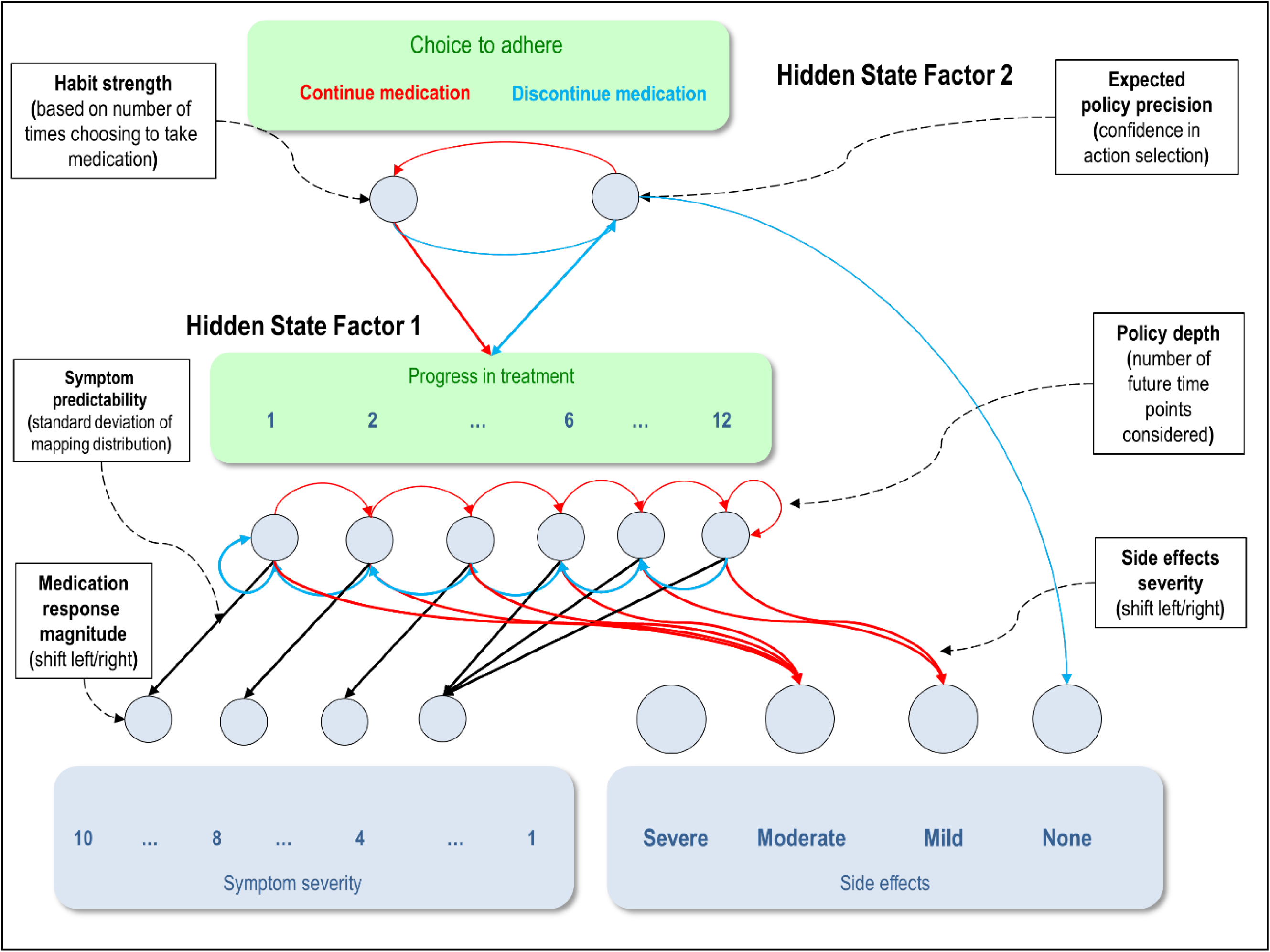
Displays the levels of hidden state factor 1 and 2 (treatment progress and adherence decisions) and their mapping to different lower-level representations of symptom levels and side effect severity (here modelled as outcomes). Each combination of levels in the two hidden state factors generated different observation patterns and state transitions with different probabilities. Red arrows correspond to the observations (and state transitions) generated when the simulated patient chooses to continue medication, whereas blue arrows correspond to the observations (and transitions) generated when the patient chooses to discontinue medication (see text for details). Black arrows correspond to observations generated by hidden states that do not depend on the selection of one policy vs. another. Each of the six parameters in the model are also illustrated with a brief description (and described in the text in more detail). For example, symptom predictability was modulated by reducing the precision of the mapping from hidden state levels to symptom severity observations. Response magnitude was modulated by shifting the observable symptom severities toward lower or higher levels. Side effect severity was modulated by shifting the observable side effect levels to higher or lower values. For example, while in the figure it shows that moderate side effects are generated in the first six weeks of treatment and mild side effects are observed thereafter, this could be shifted such that they instead transition from severe to moderate, or from mild to none. Policy depth controlled the number of future transitions in treatment progress specified for each allowable policy in the patient’s model, only allowing the patient to consider the outcomes of different policies over a limited number of weeks into the future. See the text and tables for a thorough description of these and other parameters.

Several matrices/vectors in an MDP define the probabilistic relationships between each of these variables (see figure 3A); these include matrices encoding the relationships between *states and observations* (**A-**matrices; 1 per observation modality), how *states are expected to change over time* (**B-**matrices; at least 1 per state factor – if greater than 1, each possible transition corresponds to an action option), the relative *preference for some observations* over others (**C-**matrices; 1 per observation modality), *expectations about the initial states* one will start out in (**D-**vectors; 1 per state factor), and *prior expectations about which policies one is most likely to choose* in general (**E-**vector; 1 probability entry assigned to each policy). For more mathematical detail about each of these variables/matrices, see tables 1 and 2 and figure 1.

**Figure 3.**
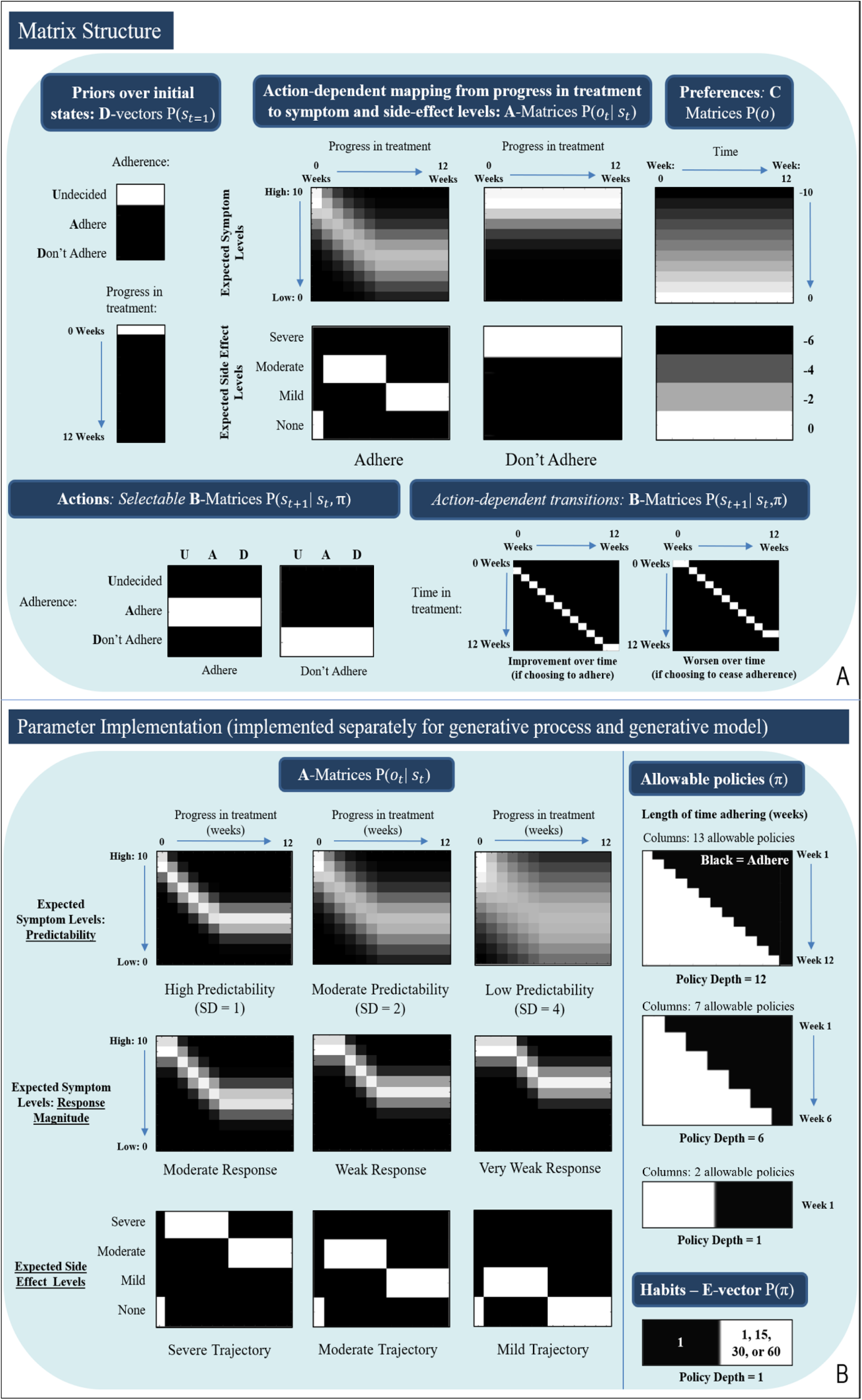
**(A)** Displays select **A**-, **B**-, **C**-matrices, and **D**-vectors, that specify the generative model. **A** encodes state-observation mappings (i.e., columns = states, rows = observations), **B** encodes policy-dependant state transitions (i.e., columns = states at time t, rows = states at time t+1), **C** encodes preference distributions (i.e., columns = time, rows = observed symptoms [top] or side effects [bottom]), and **D** encodes priors over initial states (indicating that each simulation starts in an undecided state and at 0 weeks of treatment progress), respectively. Lighter colors indicate higher probabilities. **(B)** Illustrates how **A**-matrices, the allowable policy space, and the **E**-vector (encoding priors over policies, implementing habits) are altered under different parameter values. Symptom predictability: the precision of the first **A**-matrix (i.e., encoding beliefs about how precisely different “progress in treatment” levels – corresponding to the amount of time on the medication – will lead to different observed symptom levels) could be adjusted via specifying the standard deviation of an associated Gaussian probability density function (SDs = 1, 2, and 4 are shown here from left to right). This allowed symptoms to fluctuate both upward or downward around the mean of the symptom response trajectory with different degrees of predictability. (Middle row, left) Response magnitude: Relative to that shown in figure 3A, medication response magnitude could be modulated down (from left to right) to moderate, weak, and very weak responses, corresponding to incremental shifts upward and to the right in the same matrix. (Bottom row, left) Side effect levels: The second **A**-matrix (i.e., encoding beliefs about how taking the medication will generate different side effect severities over time) could also be modulated to simulate the magnitude of side effect responses over time. From left to right, the patient either initially experienced severe side effects that eventually settled at moderate levels, initially moderate levels of that eventually settled at mild levels, or initially mild levels that eventually resolved. Each parameterization could be performed separately for the generative process and the simulated patient’s generative model – allowing the patient to observe outcomes inconsistent with her initial expectations. (Right, top) Policy depth: illustrates the allowable policies under different policy depths (number of future timesteps considered in decision-making). (Right, bottom) Habit strength: In specific simulations (with a policy depth of 1) described below, the **E**-vector (a prior over policies) was specified so as to simulate medication taking habits based on 1, 15, 30, and 60 previous choices to take medication.

The **A**-matrices were constructed such that steadily lowering (but fluctuating) symptom levels were generated for each week the patient continued to take the medication. Specifically, Gaussian distributions were specified over a symptom severity scale from 0 to 10, with empirically based means that probabilistically decreased from baseline levels over the 1^st^ 6 weeks of treatment and then remained somewhat stable thereafter (i.e., depending on the standard deviations set for these distributions). The means were based on dose response time courses characterized in a mega analysis of three SSRIs (42): 9.36, 8.04, 6.91, 5.94, 5.10, 4.38, 3.77. Adherence also generated moderate side effect levels in the first six weeks and mild side effect levels from weeks 7-12 (based on the antidepressant side effect time courses characterized in (43)). The choice to cease adherence led to the cessation of side effects and a gradual return to baseline symptom levels.

The **B**-matrices were constructed so that the patient controlled the transition from the state of ‘continue medication’ to the state of ‘discontinue medication’ at each week (i.e., if two individuals ceased taking the medication on different days during the same week, these were formally treated as the same choice). Simultaneously, each action was associated with transitions toward increasing or decreasing “progress in treatment” levels, respectively. The **C**-matrices were constructed such that the patient most preferred the lowest symptom levels (ln *P*(*o*): from − 10 to 0) and the lowest side effect severity (ln *P*(*o*): 0, −2, −4, −6). The **D**-vectors were constructed such that the patient always began in an initial state of being “undecided” about choosing to adhere, and always began in an initial state of being at 0 weeks of treatment progress. The **E**-vector was initially set such that the patient had no bias (habit) for choosing to adhere or not (i.e., a flat prior distribution over policies).

The parameters in our model are described in table 3, and their influence on the matrices/vectors defining the model are depicted in figure 3B. These correspond to individual differences in (beliefs about) the predictability and magnitude of changes in symptoms and side effects, how far into the future one considers when making decisions (policy depth), confidence in the consequences of choosing different actions (expected policy precision), and the strength of medication-taking habits.

**Table 3.** Model parameters.

| Parameter | Formal definition | Brief explanation | Observed effects |
| --- | --- | --- | --- |
| Objective symptom predictability | The standard deviation of a Gaussian probability density function specified over the **A** matrix in the <u>generative process</u> that produces observed symptom reductions over time when choosing to take the medication. | Lower levels of predictability (a high standard deviation) indicate that changes in symptoms while on medication will be less stable/predictable (around the mean trajectory) from week to week. | Generally, less predictable symptoms (greater day-to-day symptom fluctuations) deter adherence. Greater random (upward) fluctuations can also be sufficient in certain cases to change an individual's beliefs such that they now think the drug is ineffective. |
| Subjective symptom predictability | The standard deviation of a Gaussian probability density function specified over the **A** matrix encoding <u>beliefs in the generative model</u> about the symptom reductions that will be observed over time when choosing to take the medication. | Lower levels of predictability (a high standard deviation) indicate the <u>prior belief</u> that changes in symptoms while on medication will be less stable/predictable (around the mean trajectory) from week to week. | Overly optimistic beliefs about symptom predictability tend to promote adherence even in the face of highly fluctuating symptoms. |
| Objective medication response magnitude | A parameter that transforms (shifts) the **A** matrix in the <u>generative process</u> that produces observed symptom reductions over time when choosing to take the medication. This was specified as single symptom level shifts up and to the right from baseline mean symptom level trajectories (strong, moderate, weak, and very weak response magnitudes corresponded to 0, 1, 2, and 3 up/right shifts, respectively). | Lower response magnitude values indicate that there will be a longer delay before symptoms begin to decrease and that a smaller overall decrease in symptoms will be achieved by the final week of treatment. | Higher response magnitude promotes adherence because they more easily outweigh the individual's concerns about the side effects of the drug. |
| Subjective medication response magnitude | A parameter that transforms (shifts) the **A** matrix encoding <u>beliefs in the generative model</u> about the symptom reductions that will be observed over time when choosing to take the medication. This was specified as single symptom level shifts up and to the right from baseline mean symptom level trajectories (strong, moderate, weak, and very weak response magnitudes corresponded to 0, 1, 2, and 3 up/right shifts, respectively). | Lower response magnitude values indicate the <u>prior belief</u> that there will be a longer delay before symptoms begin to decrease and that a smaller overall decrease in symptoms will be achieved by the final week of treatment. | Overly optimistic beliefs about symptom response magnitudes show a trend toward improving adherence in some cases (but the effects are complex). |
| Objective side effect severity | A parameter that transforms (shifts) the **A** matrix in the <u>generative process</u> that produces observed side effects over time when choosing to take the medication. This was specified as shifts upward or downward one level in the baseline side effect time course (severe, moderate, and mild levels corresponded to shifts of 1, 0, and -1, respectively). | Lower side effect severity indicates that milder side effects will be experienced initially, and that they will eventually become less over time. | Higher side effect severity promotes non-adherence in general, as more severe side effects are not preferred. |
| Subjective side effect severity | A parameter that transforms (shifts) the **A** matrix encoding <u>beliefs in the generative model</u> about the side effects that will be observed over time when | Lower side effect severity indicates the <u>prior belief</u> that milder side effects will be experienced initially, and that they will eventually | Overly optimistic beliefs about side effects promote non-adherence, because they lead to immediate disappointment upon observing higher-than-expected side effect levels. |
|  | choosing to take the medication. This was specified as shifts upward or downward one level in the baseline side effect time course (severe, moderate, and mild levels corresponded to shifts of 1, 0, and -1, respectively). | become less over time. |  |
| Policy depth | The number of time steps over which action policies are specified. This ranged from 1 to 12 weeks, indicating the distance in the future one considers when planning how to act. A 12-week time period was chosen to match with the typical timeframe studied in antidepressant drug trials. Reducing policy depth was implemented by simply reducing the number of time steps in the simulation over which allowable policies were specified (i.e., while holding all other transition probabilities and other contingencies identical). | Lower policy depth indicates that, when people are deciding whether or not to adhere, they do not consider the changes in symptoms and side effects that will occur as far into the future. For example, they might think “I’m not going to take the medication because it’s going to make me feel bad tomorrow” without considering that it will make them feel much better after several weeks. | A high policy depth promotes adherence when an individual expects that, despite initially feeling worse (or not improving), in the long run they will ultimately feel better overall. |
| Prior on expected policy precision ($\beta$) | An “inverse temperature” parameter modulating the impact of current beliefs on policy selection. Encodes how confident an individual is in general in their ability to make the best | Lower expected policy precision (higher $\beta$ values) indicates less confidence in the choice to adhere vs. not adhere in general. When expected policy | Lower expected policy precision tends to promote adherence in the context of pessimistic beliefs about the overall effectiveness of medication. For example, an individual who does not believe the medication is likely to help may still “give the medication a try” if they have less confidence |
|  | decision regarding adherence. High precision was modeled with a value of 1, and low precision was modeled with a value of 10. | precision is low, learned medication-taking habits have a stronger influence on policy selection. | that their beliefs will lead to the best decision. |
| Habit strength | Magnitude of the prior expectation in the **E** matrix that the decision to adhere will be chosen – based on the number of times the individual has chosen to take the medication in the past. | Higher medication-taking habit strength reduces the influence of explicit beliefs about the consequences of taking medication on decision-making. | Higher medication-taking habit strength promotes adherence, even if an individual doesn't have strong beliefs that the medication will increase their quality of life overall. Adherence is chosen "out of habit" without consideration of expected effects. |

## Results

### Simulating individual differences in adherence

#### Initial simulations

The left panel of figure 4 illustrates an example simulation under one set of parameter values, in which policy depth was at its maximum value of 12, symptom predictability was moderate (SD = 2), symptom reductions were high (i.e., the simulated patient was a “strong responder” to the medication), side effects were moderate, expected policy precision was high (***β*** = 1), and no medication-taking habit had been formed (the **E**-vector distribution remained flat over all policies). Here the patient chose to adhere and observed steady symptom reductions and some initial moderate side effects that decreased to mild levels after the first six weeks. The middle panel illustrates another example simulation in which symptom predictability was moderate (SD = 2), symptom reductions were moderate, and side effects were severe. In this case, the patient chose not to adhere. In these simulations, the patient’s expectations were consistent with subsequent observations. The right panel illustrates a simulation in which the patient believed symptom predictability was higher than it really was (SD = 0.5 vs. 2) and side effects and symptom reductions were moderate (which was expected). The patient took the medication the first two weeks, but ceased adhering after observing an unexpected increase in symptom levels.

**Figure 4.**
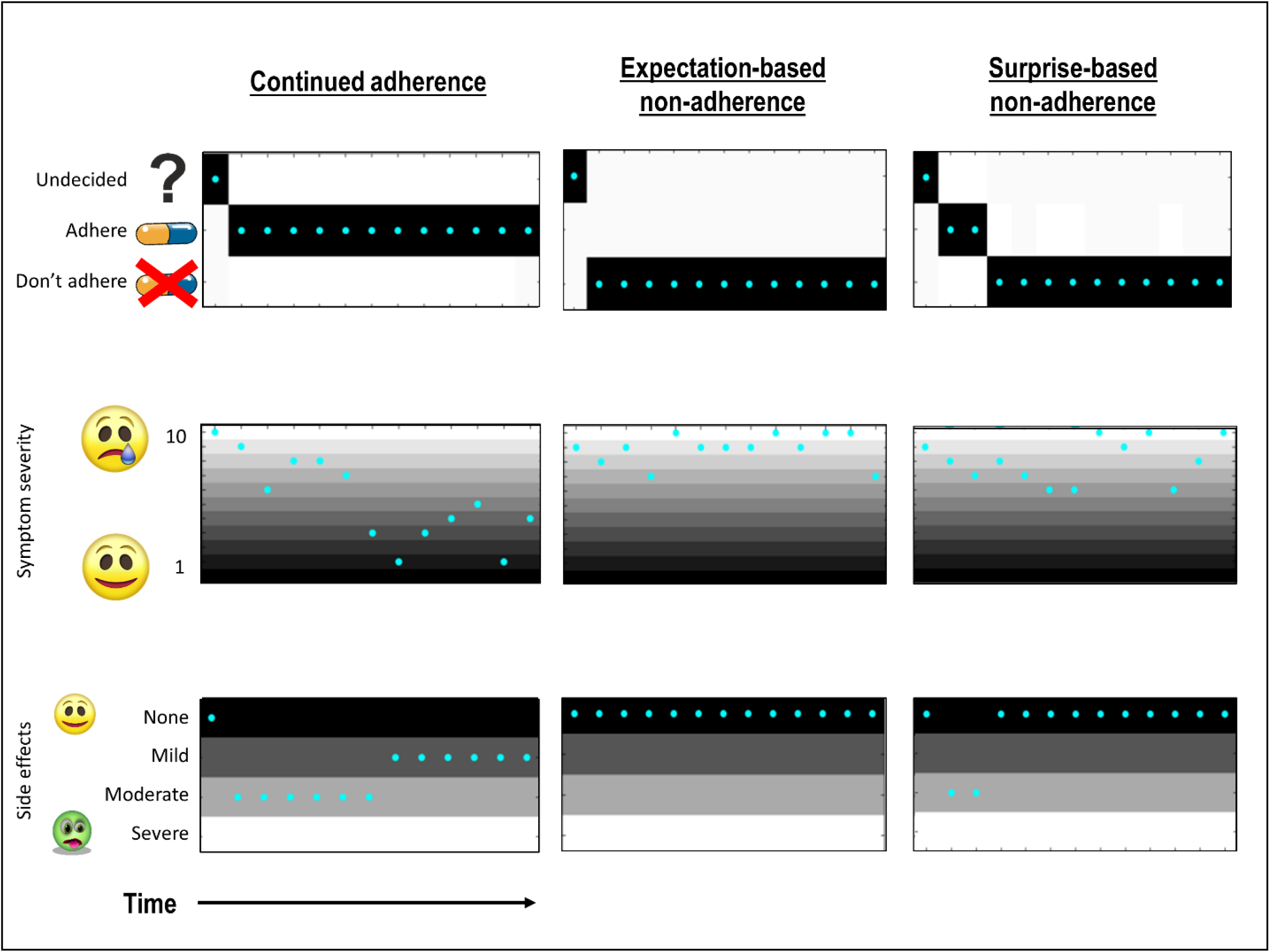
Example trials displaying different adherence decisions under different parameter values. In the top plots, cyan dots indicate the true action taken, and darker colors indicate higher levels of confidence in one action over others. In the middle and bottom plots, cyan does indicate observations, whereas darker colors indicate more strongly preferred observations. The top plots illustrate the choice to adhere or not adhere at each of the 12 weeks, the middle plots display observed symptoms over time with severities from 10 to 1, and the bottom plots illustrate observed side effects from mild to severe. Under the parameter values used in the left simulation, including (accurate) expectations for moderate side effects, strong symptom reductions, and moderate symptom predictability (SD = 2), the patient chose to adhere. Under the parameter values used in the middle simulation, in which there were (accurate) expectations for moderate symptom reductions, moderate symptom predictability (SD = 2) and severe side effects, the patient chose not to adhere – leading to an absence of side effects and symptoms continuing to fluctuate around baseline levels. Under the parameter values used in the right simulation, in which the patient expected symptoms to be more predictable than they actually were, (SD = 0.5 vs. 2) the patient initially chose to adhere but then stopped when she observed an unexpected fluctuation upward in symptom intensity and a moderate increase in side effects (side effects and symptom reductions were moderate, which was expected). After ceasing medication, side effects resolved and symptoms continued to fluctuate around baseline levels. See the main text for more details about the parameter manipulations in each simulation.

#### Parameter interactions in the context of accurate expectations

To better characterize this parameter space, we repeated the simulations above under many combinations of parameter values. Figure 5 illustrates two example plots from part of this parameter space where the patient had accurate expectations; medication response magnitude and side effect severity were fixed at moderate levels. Here, the x-axis in each plot corresponds to symptom predictability (from low to high: SDs from 4 to 0.1); the y-axis corresponds to policy depth (from 2 to 12 weeks). The left plot corresponds low expected policy precision (***β*** = 10), and the right corresponds to high expected policy precision (***β*** = 1). Black squares correspond to patients that remained adherent, whereas white squares indicate ceased adherence prior to week 12. As can be seen, there is a clear boundary in which, below a certain policy depth and level of symptom predictability, adherence ceases. Interestingly, adherence increases across the space when expected policy precision is low. This suggests that, with greater decision uncertainty (it is less clear that one policy will produce more preferred observations than others), one is more likely to adhere in an “exploratory” manner (e.g., “I don’t think this will work, but who knows?”).

**Figure 5.**
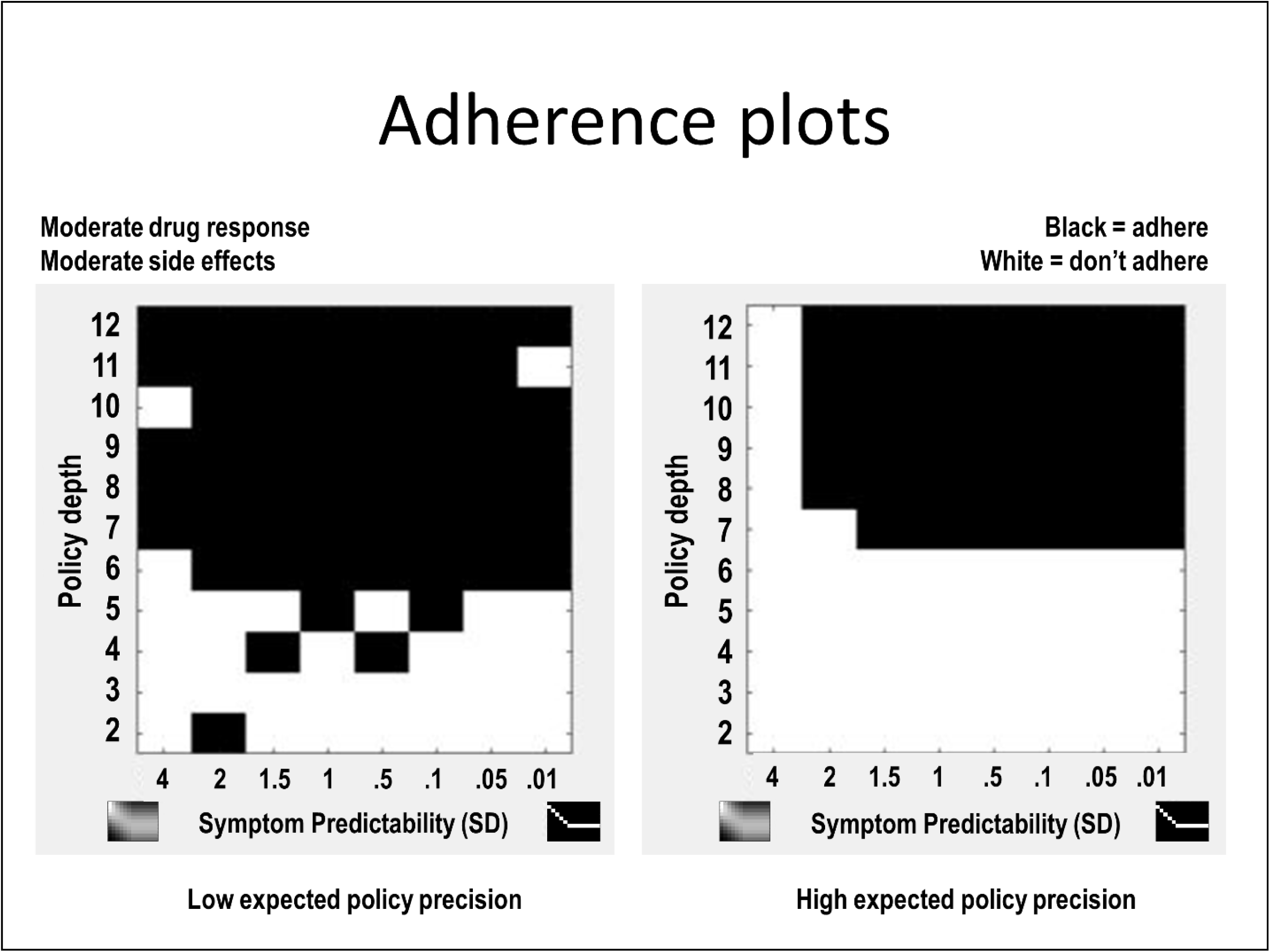
Two example plots from a larger parameter space with dimensions corresponding to different values for policy depth, symptom predictability, drug response magnitude, side effect severity, and expected policy precision. In these two plots the simulated patient had accurate expectations about what she would observe under different policies, and medication response magnitude and side effect severity were fixed at moderate levels. The x-axis in each plot corresponds to symptom predictability (from low to high: i.e., SDs from 4 to 0.1), whereas the y-axis corresponds to policy depth (from 2 to 12 weeks). The plot on the left corresponds to a simulated patient with low expected policy precision (***β*** = 10), whereas the plot on the right corresponds to a patient with high expected policy precision (***β*** = 1). Black squares in these plots correspond to patients that remained adherent all the way into week 12, whereas white squares indicate patients that ceased adherence prior to week 12. The main text for interpretation.

Figure 6 provides a more complete depiction of this parameter space under different levels of medication response magnitudes (larger x-axis across plots, from very weak to strong response) and side effect severities (larger y-axis across plots, from low to high severity). Under high side effect severity and low response magnitude, adherence does not occur no matter the policy depth or symptom predictability (upper left plot), whereas adherence occurs broadly when side effects are low and response magnitude is high (bottom left plot).

**Figure 6.**
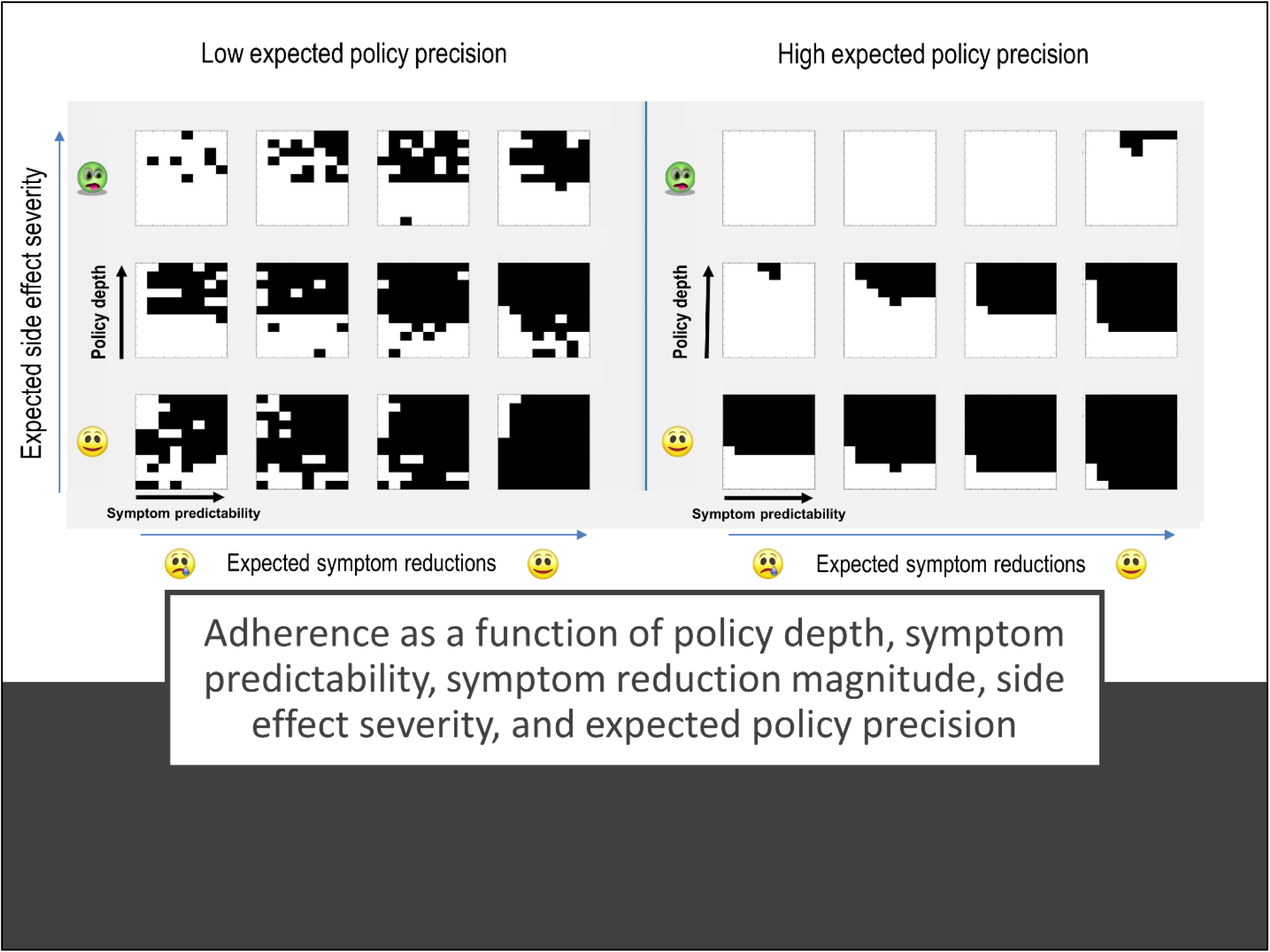
A more complete depiction of the parameter space illustrated in figure 5 including different levels of medication response magnitudes (larger x-axis across plots, left to right from very weak to strong response) and side effect severities (larger y-axis across plots, bottom to top from low to high severity). As in figure 5, within each plot black indicates continued adherence and white indicates non-adherence. Within each plot, the expected predictability of symptom changes over time increases from left to right (i.e., SDs from 4 to 0.1), and policy depth increases from bottom to top (from 2 to 12 weeks). The plot on the left corresponds to a simulated patient with low expected policy precision (***β*** = 10), whereas the plot on the right corresponds to a patient with high expected policy precision (***β*** = 1). See main text for interpretation.

#### Parameter interactions in the context of inaccurate expectations

Crucially, in the previous simulations it was assumed that the patient’s beliefs about the effects of medication were accurate. However, a patient’s beliefs need not match reality. As such, we then dissociated the patient’s beliefs from the statistics of subsequent observations to see if surprise would influence adherence in interesting ways. Supplementary figure S1 depicts some illustrative locations in this space. The x-axis in each graph corresponds to subjective beliefs about symptom predictability, where actual symptom predictability instead varies between groups of plots (see figure legend for more details). Similarly, beliefs about drug response magnitude and side effect severity are now depicted across plots within each group of plots, whereas actual drug response magnitude and side effect severity vary across groups of plots.

When observed symptom reductions were highly unreliable/noisy, the patient only remained adherent if prior expectations very precisely predicted that they would be reliable. When observed symptom reductions were instead highly consistent/reliable, the patient surprisingly also chose to adhere when expecting symptom reductions to be highly unreliable. Thorough inspections of the parameter space confirmed that, in these cases of highly reliable or unreliable symptom reductions, there was little influence of expected policy precision, drug response magnitude, or side effect severity. Other interesting results were observable in cases of objectively moderate levels of symptom reduction reliability. For example, there appears to be a general effect in which adherence was higher when expectations about side effects matched the actual side effects observed. This effect was more pronounced in the case of objectively strong drug responses; in the case of objectively weak drug responses and severe side effects, adherence levels became more dependent on the expected reliability of symptom reductions. In cases of objectively low side effects, adherence went up in general, most notably in cases of high policy depth and the belief that symptom reductions would be unreliable (unless expected side effects were severe).

The finding that adherence is higher when expected observations about side effects are confirmed appears sensible, in that, if one initially chose to “try out” adherence based on one’s expectations, non-preferred surprising observations would be more likely to “change one’s mind” later. The finding that, in the context of objectively reliable symptom reductions, adherence occurs at low but not intermediate levels of expected symptom predictability is initially more surprising. However, when symptoms are expected to fluctuate very unreliably, it makes sense retrospectively that being “pleasantly surprised” that observations are more reliable than expected would promote adherence. In contrast, if one initially expects moderate predictability, the level of surprise may not be sufficient to change one’s mind. Finally, it is fairly intuitive that, in the face of highly fluctuating symptoms, adherence would require a strong counteracting belief that they would still be reliable at future time points (e.g., “it’s been a bumpy ride to start, but I think things are going to stabilize soon”).

#### Parameter interactions in the context of habit formation

To investigate why some individuals take much longer to form strong habits than others (as reviewed in the introduction), we ran a final set of simulations in which we manipulated the strength of the patient’s habit to take medication. We manipulated the **E**-matrix to simulate the effect of a patient having previously taken the medication different numbers of times (i.e., 1, 15, 30, and 60 previous medication-taking decisions). Policy depth was set to 1, such that the patient was not forward-looking beyond the immediate expected consequences of adhering. As can be seen in figure 7, different simulated patients begin to adhere habitually after different lengths of time depending on other parameter values. Longer time periods of previous adherence lead to habitual adherence. Habits took longer to develop when expected drug responses were low and unpredictable, and when expected side effect severity was high (i.e., the medication-taking “impulse” less effectively competed against explicit planning when strongly non-preferred or unpredictable outcomes were expected to occur immediately). Low expected policy precision led to much faster habit formation. Psychologically, this might be interpreted as indicating an interesting (and somewhat paradoxical) predicted effect. That is, in the context of higher side effects and lower drug responses, individuals who are less confident in decision-making should be more likely to adhere long-term than those who confidently predict at treatment onset that the drug will not be very helpful.

**Figure 7.**
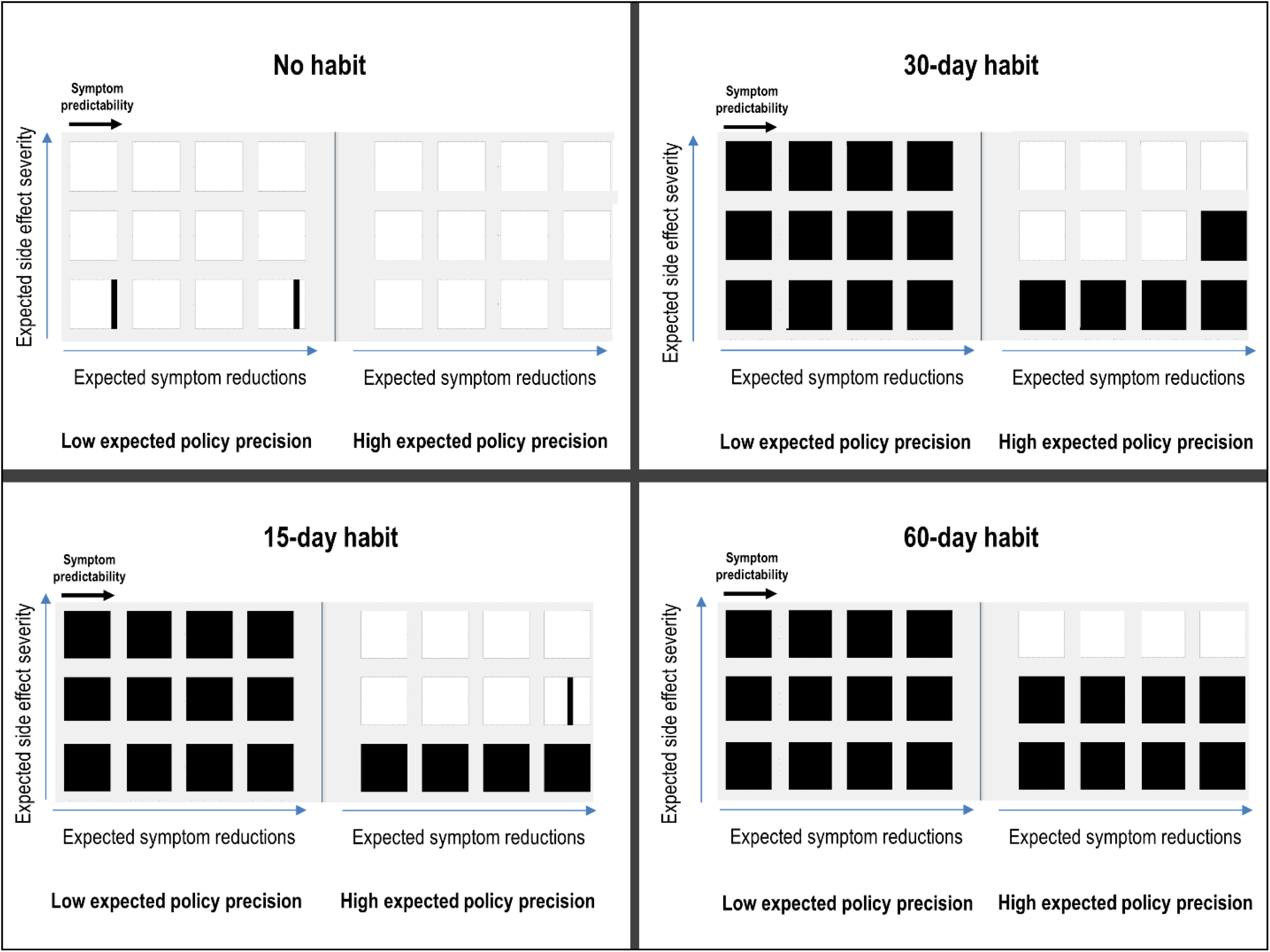
A depiction of the influence of habit formation at different habit strengths, based on the previous number of times (e.g., days) that a simulated patient has chosen to take the medication, under different values for the other parameters in the model. Within each square, black indicates adherence and white indicates non-adherence. Within each square, the expected predictability of symptom changes over time increases from left to right (i.e., SDs from 4 to 0.1). The expected magnitude of symptom reductions increases *between squares* from left to right (from very weak to strong response). The expected severity of side effects increases *between squares* from bottom to top (from low to high severity). The left and right plots within each quadrant correspond to low vs. high expected policy precision values (***β*** = 10 and ***β*** = 1, respectively). Each quadrant corresponds to the number of previous medication taking actions. Habit formation was modeled via manipulation of the **E**-vector by increasing the probability, in terms of counts (i.e., 1, 15, 30, and 60 previous medication-taking decisions), of the medication-taking action relative to other allowable actions. Here the patient also had a policy depth of 1 (i.e., y-axis within each square only has 1 value), such that she was not forward-looking beyond the immediate expected consequences of adhering. The patient otherwise had accurate expectations about the observations she would make under different actions. See main text for interpretation.

## Discussion

The goal of this manuscript was to provide a computational framework to develop novel approaches that can be used to increase adherence. Simulations were used to quantitatively illustrate distinct decision-making processes that could contribute to differences in adherence behavior. These simulations demonstrate how adherence can be influenced by several underlying computational processes. This supports the plausibility of heterogeneous causes of non-adherence (for a summary of possible causes in our model, see table 3). For example, while some patients may focus on proximal as opposed to distal future outcomes (i.e., low policy depth), others may focus on distal future outcomes but believe those outcomes are highly unreliable or that they will be worse overall if they follow treatment recommendations. Some patients may be characterized by intermediate combinations, such as intermediate policy depth and competing beliefs about moderately beneficial medication effects and moderately aversive side effects, both weighted by the relative confidence they have in each of those beliefs. Finally, differences in previous experience taking medication can lead to differences in adherence-promoting habit formation – where each of the other factors described above can influence how quickly such habits gain sufficient strength to maintain long-term adherence behavior. One advantage of the model is that each of these factors and their interactions can be simulated mathematically. The pragmatic utility of this approach will depend on developing experimental paradigms and/or questionnaires to estimate parameter values based on a given individual’s behavior. This model could potentially make predictions about where interesting “tipping points” would begin to favor non-adherence over adherence in that individual’s decision process.

At least three important research implications follow. First, it will be important to either identify or design simple measures (e.g., that could be administered in a clinic) capable of characterizing where an individual patient falls within the parameter space we have described. Such measures could potentially improve prediction of who will and will not adhere to a prescribed treatment. For example, our model would predict lower adherence in those who self-report overconfidence in their current plan of action, the tendency to focus on short-term outcomes, and either pessimistic or unrealistic expectations about how their symptoms and side effects would change over time.

With respect to existing measures, the values for some parameters might be assessed by those used within the “necessity versus concerns” framework, such as the Beliefs about Medicines Questionnaire (12) or the revised Medication Adherence Reasons Scale (50) – perhaps most plausibly the parameters associated with beliefs about the magnitude of symptom reductions and side effects. In table 4, we have also listed a number of example self-report items that, based on our model (and straightforward extensions of it), could be useful for gathering information about a patient’s adherence-relevant beliefs. Once in a validated form, this type of questionnaire could be tested for its utility in predicting adherence behavior in advance.

**Table 4:**
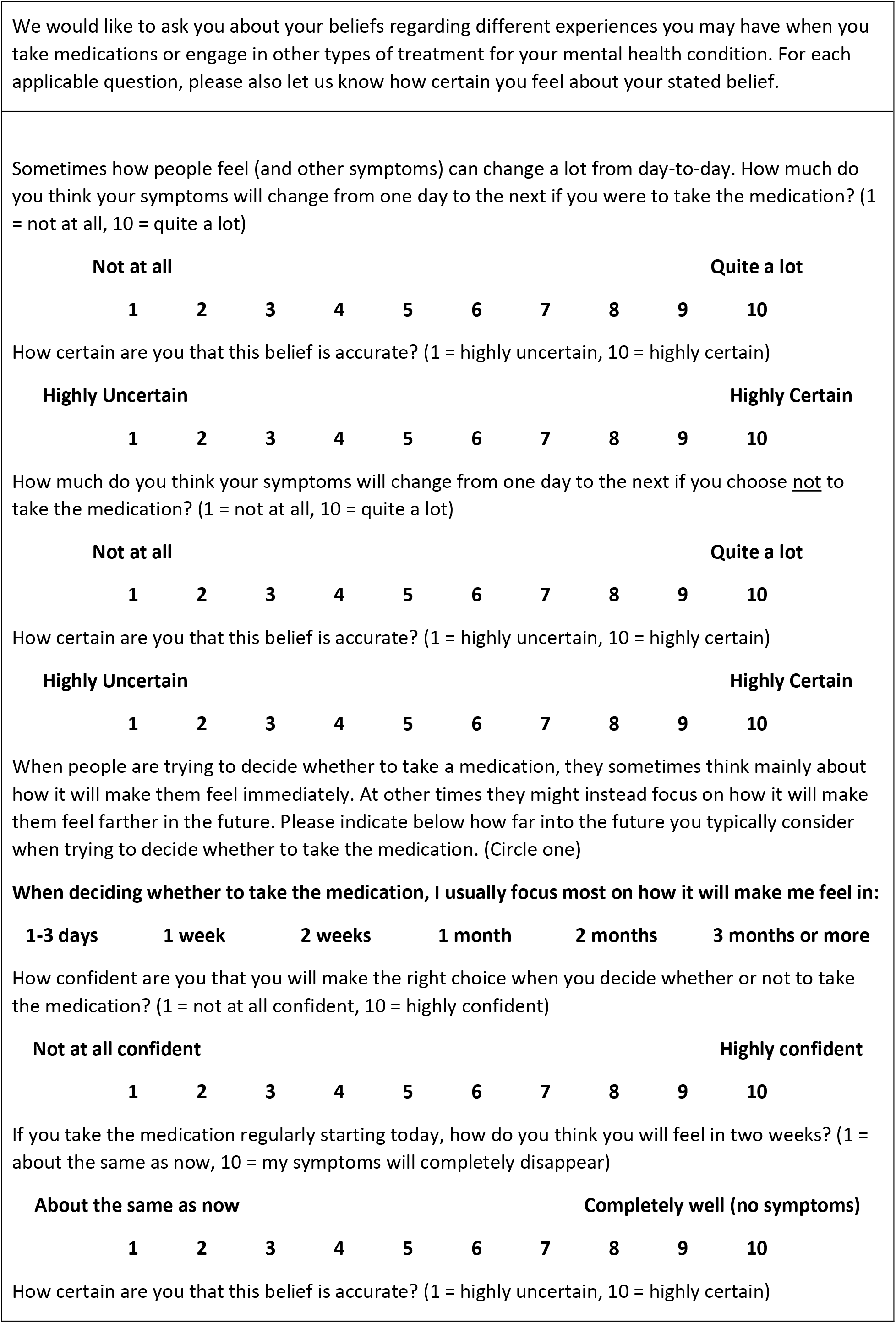
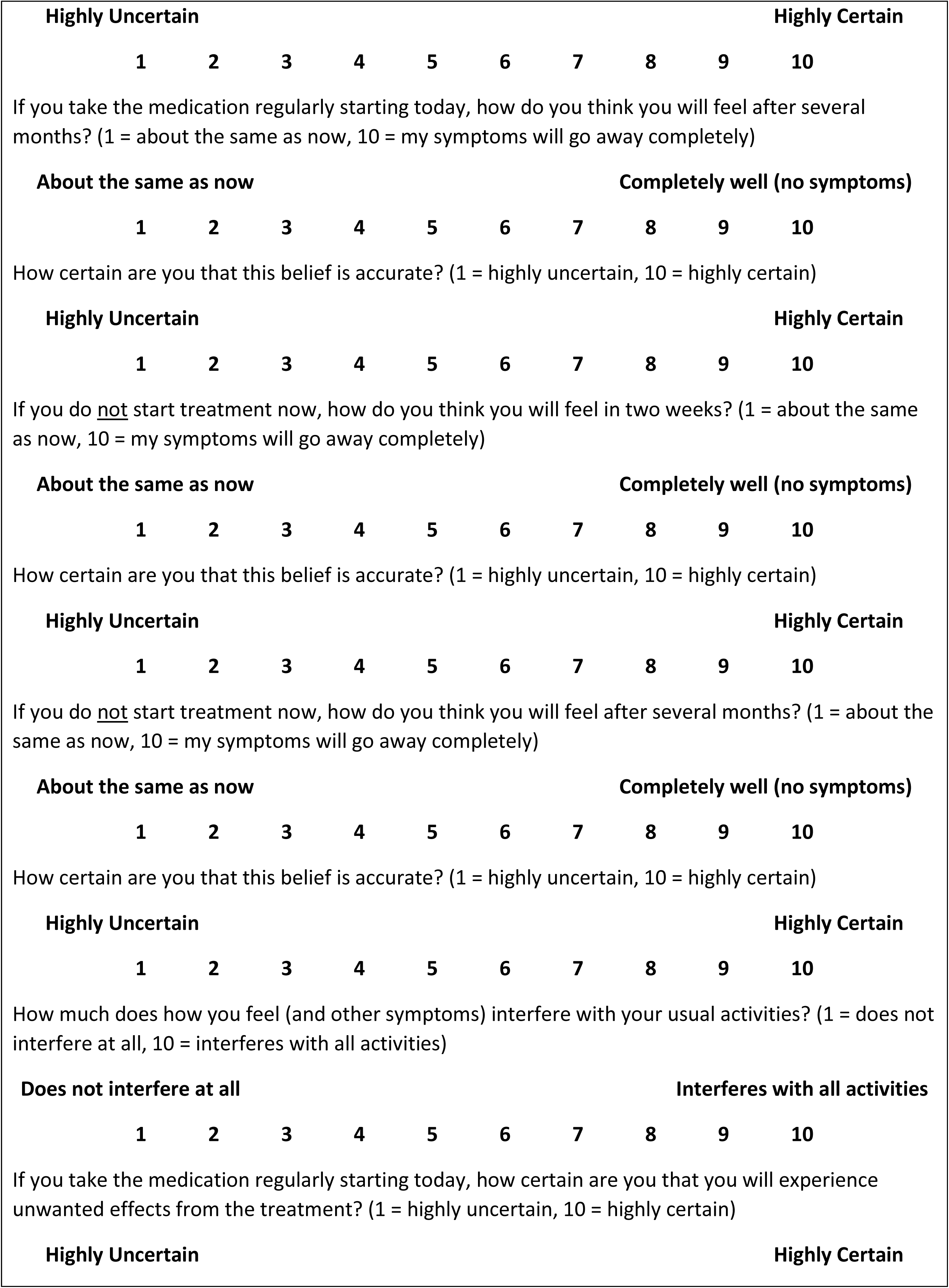
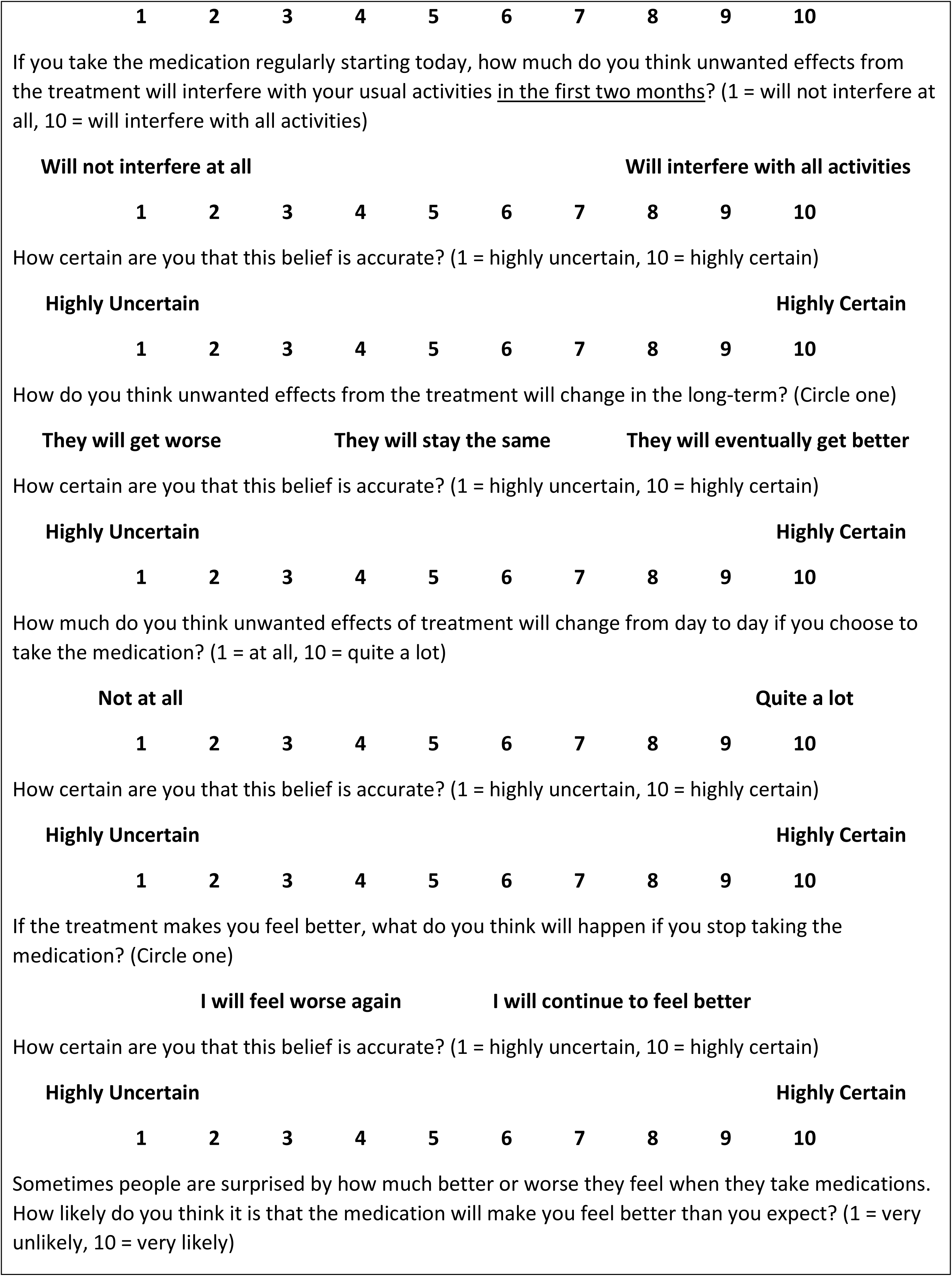
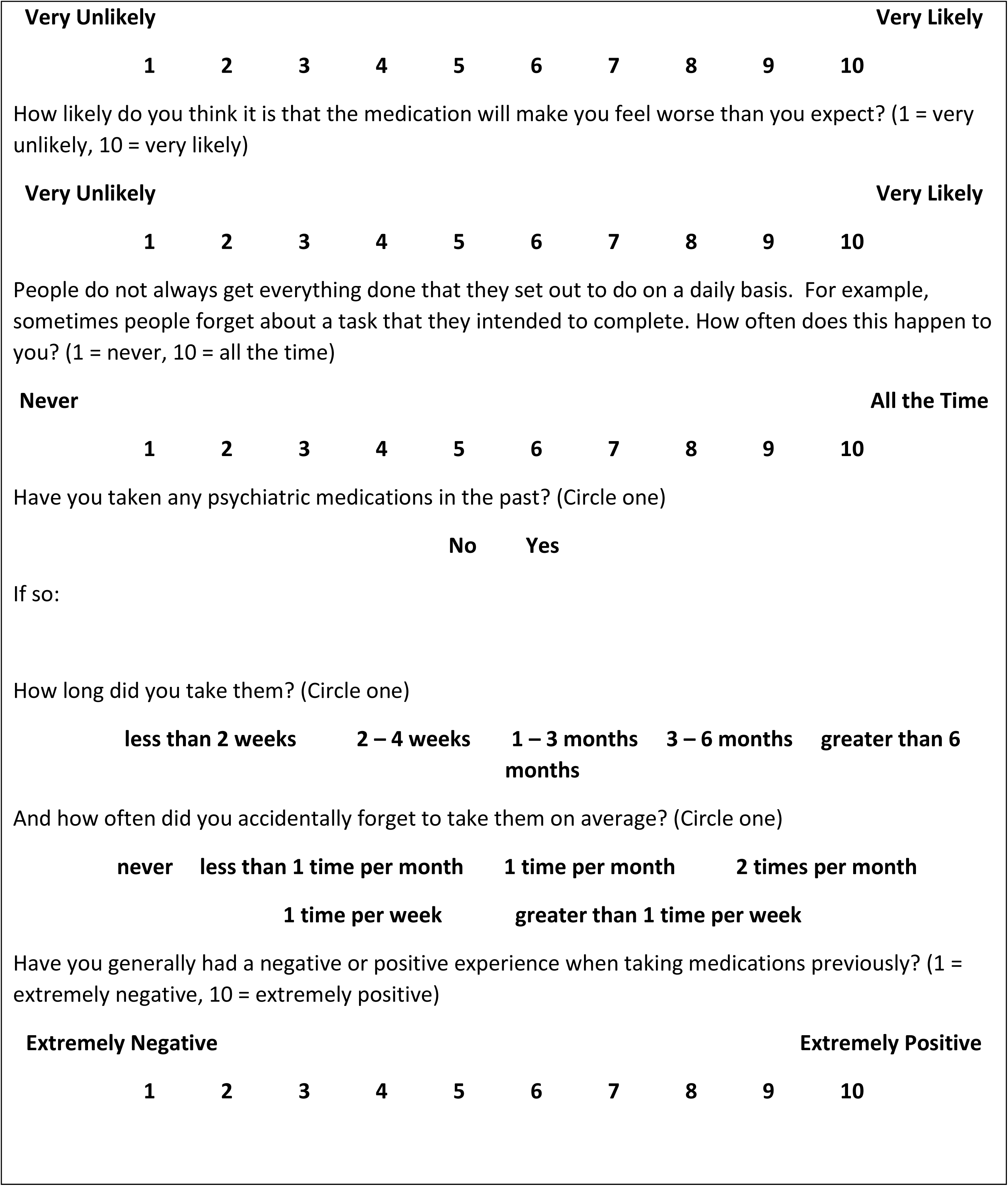
Assessment questions motivated by our model:

The values of other parameters, in contrast, might be informed by personality measures. For example, the behavior associated with persistence (20) and locus of control (26) could follow from a combination of beliefs in high future predictability and high expected policy precision (and likely preferences with strong magnitudes as well). Low self-efficacy (21) could instead follow from a combination of low expected policy precision and precise expectations for non-preferred outcomes (while optimism may correspond to precise expectations for preferred outcomes, perhaps with or without taking medication – which could explain why more optimistic individuals are less likely to adhere; (24)). This is speculative, however, and will need to be examined in future work.

A second implication is that it will be important within the field of computational psychiatry to attempt to develop tasks, or find ways of acquiring detailed adherence behavior data, that could be used in conjunction with a formal model (such as the one we described above) to explicitly fit to patient behavior. This would potentially allow for a more detailed and informative way to phenotype decision-making processes within individual patients and afford more precise empirical predictions. It should be mentioned, however, that this may be challenging since the behavior in question involves simple binary decisions. If multiple parameter value combinations can produce the exact same behavior, this prevents identification of unique parameter values that best explain individual patients’ behavior. As such, this endeavor will likely require use of very detailed behavioral data, perhaps involving day-by-day medication-taking actions.

A third implication pertains to the need for effective adherence-promoting interventions. As reviewed above, current interventions have met with limited success (36), which could be due to heterogeneity in underlying mechanisms as well as a failure to specifically target those mechanisms. Based on the factors highlighted in our model, it might be possible to improve the ability of current interventions (or design new interventions) to intervene in a targeted manner. The degree of modifiability in those factors is an open question, but certainly one worth pursuing. Based on our simulations, we would predict that both of the following should improve adherence:

1. Increasing future-oriented thinking (policy depth)
2. Attenuating overconfidence (expected policy precision)

Additionally, we would predict that adherence would be improved by first providing patients with available information about what to expect regarding likely symptom and side effect trajectories (and the actual uncertainty around those expectations) – and then using observations early in treatment to update expectations about the most likely trajectory for that individual. This goes beyond simple psychoeducation, which alone has been found to be ineffective in improving antidepressant adherence rates (51), and should motivate future research into how individual symptom and side effect trajectories could be diagnosed as early as possible. This also relates to potential uses of our model for psychoeducation. For example, by making the various model components salient to a patient (e.g., by considering their own answers to the example questions in table 4), they could become more aware of how these beliefs influence their own decision-making.

It is important to stress the oversimplified and incomplete nature of the model we have presented. For simulations one must unavoidably make somewhat arbitrary decisions about the values that should be assigned to fixed parameters (and what parameters to fix). For example, we chose to set the preference magnitudes for side effects and symptom levels to specific values, and the simulation results would be expected to differ somewhat if different values had been chosen. A possible future direction could be to calibrate the model to individuals’ preference magnitudes (e.g., how much they personally dislike particular symptoms and side effects) and use this to simulate/predict their future adherence behavior. We also did not manipulate beliefs about the predictability of side effects, which could influence adherence decisions. In these cases, we instead chose to hold these factors fixed and examine the effect of altering the dynamics of observed symptoms/side effect levels over time on versus off treatment.

There may also be additional factors that were not modeled explicitly, but that could be simulated within the model we have presented (and in principle used in characterizing patients’ decisions). For example, some individuals may simply forget to take their medication a couple of times and then cease altogether (50). In our model, this could be captured in part by low expected policy precision (i.e., increasing randomness in behavior), but would need to be combined with other factors such as the belief that all progress has been lost (as could be encoded within the individual’s transition beliefs [**B**-matrix]). As another example, some people might cease medication after experiencing symptom improvement because they expect such improvements to remain stable after ceasing to take the drug (and the belief that side effects would go away (52)). This could also be captured in our model with straightforward adjustments to the **B**-matrix.

## Conclusion

This active inference model is an important first step in developing a precise and quantitative delineation of decision-making factors and dynamics that influence a patient’s decision to adhere to treatment. Because of the model’s generality, it can also be very easily extended to model adherence to other medications, simply by inserting the symptom reduction and side effect profiles that characterize those medications. For example, it follows from the general model structure that adherence to immediately rewarding medications (e.g. benzodiazepines) would be high and promote fast habit formation, but behavior would also be influenced by the same parameters used here to investigate antidepressant adherence. The next steps in using these models will require identifying means of empirically characterizing and intervening on these mechanisms in an individualized manner.

## Supporting information

adherence_model.txt

## Software note

Although the generative model – specified by the various matrices described in this paper – changes from application to application, the belief updates are generic and can be implemented using standard routines (here **spm_MDP_VB_X.m**). These routines are available as Matlab code in the DEM toolbox of the most recent versions of SPM academic software: http://www.fil.ion.ucl.ac.uk/spm/. The simulations in this paper can be reproduced via running the Matlab code included here as supplementary material (**adherence_model.m**).

## Acknowledgments

This work has been supported in part by The William K. Warren Foundation, the National Institute of Mental Health Award Numbers K23MH112949 (SSK), and the National Institute of General Medical Sciences Center Grant Award Number 1P20GM121312 (MPP and SSK). The content is solely the responsibility of the authors and does not necessarily represent the official views of the National Institutes of Health. The authors would like to acknowledge Michael Moutoussis for his helpful suggestions during revision of the manuscript.

## References

1. Aikens JE, Klinkman MS (2012): Changes in patients’ beliefs about their antidepressant during the acute phase of depression treatment. General hospital psychiatry. 34:221–226.

2. Bosworth HB, Granger BB, Mendys P, Brindis R, Burkholder R, Czajkowski SM, et al. (2011): Medication adherence: a call for action. American heart journal. 162:412–424.

3. Serna MC, Cruz I, Real J, Gasco E, Galvan L (2010): Duration and adherence of antidepressant treatment (2003 to 2007) based on prescription database. European psychiatry : the journal of the Association of European Psychiatrists. 25:206–213.

4. Olfson M, Marcus SC, Tedeschi M, Wan GJ (2006): Continuity of antidepressant treatment for adults with depression in the United States. The American journal of psychiatry. 163:101–108.

5. Kim KH, Lee SM, Paik JW, Kim NS (2011): The effects of continuous antidepressant treatment during the first 6 months on relapse or recurrence of depression. Journal of affective disorders. 132:121–129.

6. Yue Z, Cai C, Ai-Fang Y, Feng-Min T, Li C, Bin W (2014): The effect of placebo adherence on reducing cardiovascular mortality: a meta-analysis. Clinical research in cardiology : official journal of the German Cardiac Society. 103:229–235.

7. Simpson SH, Eurich DT, Majumdar SR, Padwal RS, Tsuyuki RT, Varney J, et al. (2006): A meta-analysis of the association between adherence to drug therapy and mortality. BMJ (Clinical research ed). 333:15.

8. Sofi F, Cesari F, Abbate R, Gensini GF, Casini A (2008): Adherence to Mediterranean diet and health status: meta-analysis. BMJ (Clinical research ed). 337:a1344.

9. Gibbons RD, Hur K, Bhaumik DK, Mann JJ (2005): The relationship between antidepressant medication use and rate of suicide. Archives of general psychiatry. 62:165–172.

10. Beatty L, Binnion C (2016): A Systematic Review of Predictors of, and Reasons for, Adherence to Online Psychological Interventions. International journal of behavioral medicine. 23:776–794.

11. Murata A, Kanbayashi T, Shimizu T, Miura M (2012): Risk factors for drug nonadherence in antidepressant-treated patients and implications of pharmacist adherence instructions for adherence improvement. Patient preference and adherence. 6:863–869.

12. Horne R, Weinman J, Hankins M (1999): The beliefs about medicines questionnaire: The development and evaluation of a new method for assessing the cognitive representation of medication. Psychology & Health. 14:1–24.

13. De las Cuevas C, Penate W, Sanz EJ (2014): Risk factors for non-adherence to antidepressant treatment in patients with mood disorders. European journal of clinical pharmacology. 70:89–98.

14. Burnett-Zeigler I, Kim HM, Chiang C, Kavanagh J, Zivin K, Rockefeller K, et al. (2014): The association between race and gender, treatment attitudes, and antidepressant treatment adherence. International journal of geriatric psychiatry. 29:169–177.

15. Stetler C (2014): Adherence, expectations and the placebo response: why is good adherence to an inert treatment beneficial? Psychol Health. 29:127–140.

16. van Geffen EC, Heerdink ER, Hugtenburg JG, Siero FW, Egberts AC, van Hulten R (2010): Patients’ perceptions and illness severity at start of antidepressant treatment in general practice. The International journal of pharmacy practice. 18:217–225.

17. Phillips LA, Diefenbach MA, Kronish IM, Negron RM, Horowitz CR (2014): The necessity-concerns framework: a multidimensional theory benefits from multidimensional analysis. Annals of behavioral medicine : a publication of the Society of Behavioral Medicine. 48:7–16.

18. Foot H, La Caze A, Gujral G, Cottrell N (2016): The necessity-concerns framework predicts adherence to medication in multiple illness conditions: A meta-analysis. Patient education and counseling. 99:706–717.

19. Cloninger CR, Przybeck TR, Svrakic DM (1991): The Tridimensional Personality Questionnaire: U.S. normative data. PsycholRep. 69:1047–1057.

20. Cloninger CR, Zohar AH, Hirschmann S, Dahan D (2012): The psychological costs and benefits of being highly persistent: personality profiles distinguish mood disorders from anxiety disorders. Journal of affective disorders. 136:758–766.

21. Judge TA, Erez A, Bono JE, Thoresen CJ (2002): Are measures of self-esteem, neuroticism, locus of control, and generalized self-efficacy indicators of a common core construct? Journal of personality and social psychology. 83:693–710.

22. Burra TA, Chen E, McIntyre RS, Grace SL, Blackmore ER, Stewart DE (2007): Predictors of self-reported antidepressant adherence. Behavioral medicine (Washington, DC). 32:127–134.

23. Vangeli E, Bakhshi S, Baker A, Fisher A, Bucknor D, Mrowietz U, et al. (2015): A Systematic Review of Factors Associated with Non-Adherence to Treatment for Immune-Mediated Inflammatory Diseases. Advances in therapy. 32:983–1028.

24. Kronstrom K, Karlsson H, Nabi H, Oksanen T, Salo P, Sjosten N, et al. (2014): Optimism and pessimism as predictors of initiating and ending an antidepressant medication treatment. Nordic journal of psychiatry. 68:1–7.

25. Duckworth AL, Kern ML (2011): A Meta-Analysis of the Convergent Validity of Self-Control Measures. Journal of research in personality. 45:259–268.

26. Voils CI, Steffens DC, Flint EP, Bosworth HB (2005): Social support and locus of control as predictors of adherence to antidepressant medication in an elderly population. The American journal of geriatric psychiatry : official journal of the American Association for Geriatric Psychiatry. 13:157–165.

27. Hong TB, Oddone EZ, Dudley TK, Bosworth HB (2006): Medication barriers and anti-hypertensive medication adherence: the moderating role of locus of control. Psychology, health & medicine. 11:20–28.

28. Gardner B, de Bruijn GJ, Lally P (2011): A systematic review and meta-analysis of applications of the Self-Report Habit Index to nutrition and physical activity behaviours. Annals of behavioral medicine : a publication of the Society of Behavioral Medicine. 42:174–187.

29. Bolman C, Arwert TG, Vollink T (2011): Adherence to prophylactic asthma medication: habit strength and cognitions. Heart & lung : the journal of critical care. 40:63–75.

30. Kothe EJ, Sainsbury K, Smith L, Mullan BA (2015): Explaining the intention-behaviour gap in gluten-free diet adherence: The moderating roles of habit and perceived behavioural control. Journal of health psychology. 20:580–591.

31. Lally P, van Jaarsveld CHM, Potts HWW, Wardle J (2010): How are habits formed: Modelling habit formation in the real world. European Journal of Social Psychology. 40:998–1009.

32. Vergouwen AC, Bakker A, Katon WJ, Verheij TJ, Koerselman F (2003): Improving adherence to antidepressants: a systematic review of interventions. The Journal of clinical psychiatry. 64:1415–1420.

33. Akerblad AC, Bengtsson F, Ekselius L, von Knorring L (2003): Effects of an educational compliance enhancement programme and therapeutic drug monitoring on treatment adherence in depressed patients managed by general practitioners. International clinical psychopharmacology. 18:347–354.

34. Peveler R, George C, Kinmonth AL, Campbell M, Thompson C (1999): Effect of antidepressant drug counselling and information leaflets on adherence to drug treatment in primary care: randomised controlled trial. BMJ (Clinical research ed). 319:612–615.

35. Brook OH, van Hout H, Stalman W, Nieuwenhuyse H, Bakker B, Heerdink E, et al. (2005): A pharmacy-based coaching program to improve adherence to antidepressant treatment among primary care patients. Psychiatric services (Washington, DC). 56:487–489.

36. Nieuwlaat R, Wilczynski N, Navarro T, Hobson N, Jeffery R, Keepanasseril A, et al. (2014): Interventions for enhancing medication adherence. The Cochrane database of systematic reviews.CD000011.

37. Friston K, Stephan K, Montague R, Dolan R (2014): Computational psychiatry: the brain as a phantastic organ. The lancet Psychiatry. 1:148–158.

38. Huys Q, Maia T, Frank M (2016): Computational psychiatry as a bridge from neuroscience to clinical applications. Nature Neuroscience. 19:404–413.

39. Montague P, Dolan R, Friston K, Dayan P (2012): Computational psychiatry. Trends in Cognitive Sciences. 16:72–80.

40. Petzschner F, Weber L, Gard T, Stephan K (2017): Computational Psychosomatics and Computational Psychiatry: Toward a Joint Framework for Differential Diagnosis. Biological Psychiatry. 82:421–430.

41. Friston K, FitzGerald T, Rigoli F, Schwartenbeck P, Pezzulo G (2017): Active Inference: A Process Theory. Neural Computation. 29:1–49.

42. Hieronymus F, Nilsson S, Eriksson E (2016): A mega-analysis of fixed-dose trials reveals dose-dependency and a rapid onset of action for the antidepressant effect of three selective serotonin reuptake inhibitors. Transl Psychiatry. 6:e834.

43. Crawford AA, Lewis S, Nutt D, Peters TJ, Cowen P, O’Donovan MC, et al. (2014): Adverse effects from antidepressant treatment: randomised controlled trial of 601 depressed individuals. Psychopharmacology (Berl). 231:2921–2931.

44. Doshi F, Pineau J, Roy N (2008): Reinforcement Learning with Limited Reinforcement: Using Bayes Risk for Active Learning in POMDPs. Proc Int Conf Mach Learn. 301:256–263.

45. Kaelbling LP, Littman ML, Cassandra AR (1998): Planning and acting in partially observable stochastic domains. Artificial Intelligence. 101:99–134.

46. Friston K, Parr T, de Vries B (2017): The graphical brain: Belief propagation and active inference. Network Neuroscience. 1:381–414.

47. Friston K, Rosch R, Parr T, Price C, Bowman H (2018): Deep temporal models and active inference. Neuroscience & Biobehavioral Reviews. 90:486–501.

48. Parr T, Friston K (2018): The Discrete and Continuous Brain: From Decisions to Movement—and Back Again. Neural Computation.1–29.

49. Parr T, Markovic D, Kiebel S, Friston K (2019): Neuronal message passing using Mean-field, Bethe, and Marginal approximations. Scientific Reports. 9:1889.

50. Unni EJ, Olson JL, Farris KB (2014): Revision and validation of Medication Adherence Reasons Scale (MAR-Scale). Current medical research and opinion. 30:211–221.

51. Chong WW, Aslani P, Chen TF (2011): Effectiveness of interventions to improve antidepressant medication adherence: a systematic review. Int J Clin Pract. 65:954–975.

52. Shelton RC (2001): Steps Following Attainment of Remission: Discontinuation of Antidepressant Therapy. Prim Care Companion J Clin Psychiatry. 3:168–174.

53. Friston K, Lin M, Frith C, Pezzulo G, Hobson J, Ondobaka S (2017): Active Inference, Curiosity and Insight. Neural Computation. 29:2633–2683.

