## Supplementary material for "An Active Inference Approach to Dissecting Reasons for Non-Adherence to Antidepressants": Figure S1

***Supplemental Information***

**Figure S1.** A depiction of several locations within a higher dimensional parameter space than that shown in figures 5 and 6, in which the patient’s beliefs about symptom predictability, drug response magnitude, and side effect severity could differ from the true values generating their observations. Here the x-axis within each plot corresponds to the patient’s beliefs about symptom predictability while the y-axis within each plot still corresponds to policy depth. The larger x-and y-axes across each group of plots now corresponds to the patient’s beliefs about drug response magnitude and side effect severity, respectively. Each group of plots in turn corresponds to different combinations of the actual parameter values generating the patient’s observations. (A) The top and bottom plots in panel A illustrate the influence of very low (SD = 4) vs. very high (SD = 0.01) symptom predictability. (B) The 4 plots in panel B instead illustrate the influence of different combinations of objectively high/low side effect severities and drug response magnitudes under cases of moderately unreliable symptom reductions (SD = 0.5). See main text for interpretations.

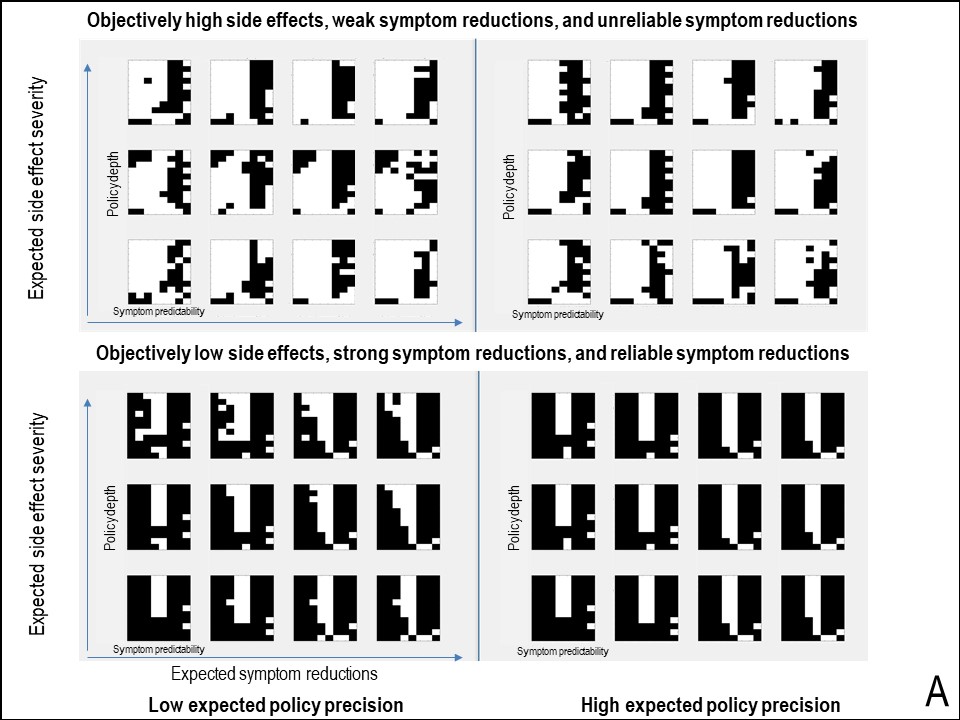

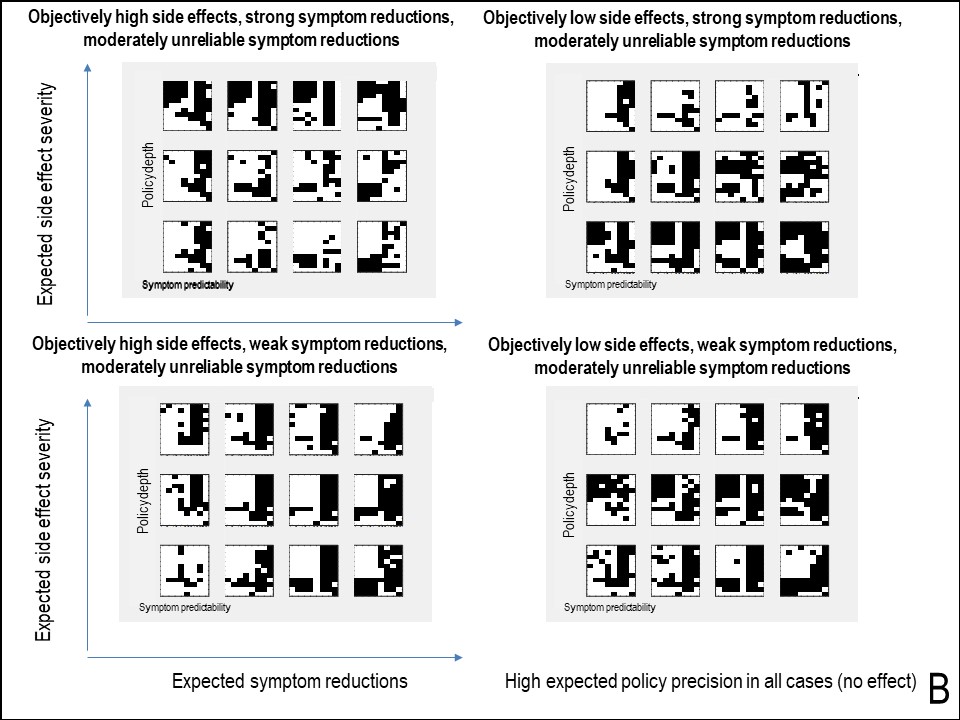
